# Predicting changes in protein thermodynamic stability upon point mutation with deep 3D convolutional neural networks

**DOI:** 10.1101/2020.02.28.959874

**Authors:** Bian Li, Yucheng T. Yang, John A. Capra, Mark B. Gerstein

**Affiliations:** Program in Computational Biology and Bioinformatics, Yale University, New Haven, CT 06520, USA; Department of Molecular Biophysics and Biochemistry, Yale University, New Haven, CT 06520, USA; Department of Computer Science, Yale University, New Haven, CT 06520, USA; Department of Biological Sciences and Vanderbilt Genetics Institute, Vanderbilt University, Nashville, TN 37235, USA

## Abstract

Predicting mutation-induced changes in protein thermodynamic stability (∆∆G) is of great interest in protein engineering, variant interpretation, and understanding protein biophysics. We introduce ThermoNet, a deep, 3D-convolutional neural network designed for structure-based prediction of ∆∆Gs upon point mutation. To leverage the image-processing power inherent in convolutional neural networks, we treat protein structures as if they were multi-channel 3D images. In particular, the inputs to ThermoNet are uniformly constructed as multi-channel voxel grids based on biophysical properties derived from raw atom coordinates. We train and evaluate ThermoNet with a curated data set that accounts for protein homology and is balanced with direct and reverse mutations; this provides a framework for addressing biases that have likely influenced many previous ∆∆G prediction methods. ThermoNet demonstrates performance comparable to the best available methods on the widely used S^sym^ test set. However, ThermoNet accurately predicts the effects of both stabilizing and destabilizing mutations, while most other methods exhibit a strong bias towards predicting destabilization. We further show that homology between S^sym^ and widely used training sets like S2648 and VariBench has likely led to overestimated performance in previous studies. Finally, we demonstrate the practical utility of ThermoNet in predicting the ∆∆Gs for two clinically relevant proteins, p53 and myoglobin, and for pathogenic and benign missense variants from ClinVar. Overall, our results suggest that 3D convolutional neural networks can model the complex, non-linear interactions perturbed by mutations, directly from biophysical properties of atoms.

**Author Summary:** The thermodynamic stability of a protein, usually represented as the Gibbs free energy for the biophysical process of protein folding (∆G), is a fundamental thermodynamic quantity. Predicting mutation-induced changes in protein thermodynamic stability (∆∆G) is of great interest in protein engineering, variant interpretation, and understanding protein biophysics. However, predicting ∆∆Gs in an accurate and unbiased manner has been a long-standing challenge in the field of computational biology. In this work, we introduce ThermoNet, a deep, 3D-convolutional neural network designed for structure-based ∆∆G prediction. To leverage the image-processing power inherent in convolutional neural networks, we treat protein structures as if they were multi-channel 3D images. ThermoNet demonstrates performance comparable to the best available methods. However, ThermoNet accurately predicts the effects of both stabilizing and destabilizing mutations, while most other methods exhibit a strong bias towards predicting destabilization. We also demonstrate that the presence of homologous proteins in commonly used training and testing sets for ∆∆G prediction methods has likely influenced previous performance estimates. Finally, we highlight the practical utility of ThermoNet by applying it to predicting the ∆∆Gs for two clinically relevant proteins, p53 and myoglobin, and for pathogenic and benign missense variants from ClinVar.

## Introduction

The thermodynamic stability of a protein, usually represented as the Gibbs free energy for the biophysical process of protein folding (∆G), is a fundamental thermodynamic quantity. The magnitude of ∆G is collectively determined by the intramolecular interactions between amino acid residues within the protein and the interactions between the protein and the physiological environment around it (1). When a mutation causes amino acid substitution in a protein, it is likely that the stability of the mutant protein will be affected compared to the wild type. (Note that the term “wild type” is not preferred because humans have substantial protein-coding genetic diversity (2). A better term would be a “reference state”. However, as it will not cause confusion in this work and for consistency with previous work, we use “wild type” throughout the text.) This change in protein thermodynamic stability (i.e. ∆∆G) incurred by mutation is of fundamental importance to medicine and biotechnology. Many disease-causing mutations are single-point amino acid substitutions that lead to a substantial ∆∆G of the corresponding protein, and such single-point mutations are a key mechanism underlying a wide spectrum of molecular disorders (3–5). Given the huge number of variants discovered by large-scale population-level exome and genome sequencing studies and clinical genetic tests, there is a tremendous interest in predicting whether these variants are likely to exert any impact on protein function. In addition, in developing new biopharmaceuticals, one of the early goals is usually to design proteins with the intended thermodynamic stability. However, this task is often laborious, if not impossible, and usually involves experimentally screening an enormous number of mutant proteins (6). Thus, it is desirable to have an efficient and accurate computational tool to prioritize the set of mutant proteins to be experimentally tested.

Toward these goals, several programs have been developed for estimating ∆∆Gs. These methods either rely on explicit biophysical modeling of amino acid interactions coupled with conformational sampling of protein structures (7–11) or apply machine/statistical learning to extract patterns from various types of amino acid sequence, evolutionary, and/or protein structural features (12–20). While these methods have been useful in many applications (21, 22), they have substantial limitations. For example, physics-based methods are computationally demanding and low-throughput; these challenges have largely prevented them from being applied to large-scale protein engineering and variant interpretation tasks. On the other hand, several studies have highlighted significant bias in the predictions of machine learning-based methods; they tend to predict mutations as destabilizing more often than stabilizing (19, 23–25). The main source of this bias likely comes from the fact that the training sets are dominated by experiment-derived destabilizing mutations and that machine learning methods are prone to overfitting to training sets (24, 26). Thus, there is a need for new methods that can make quantitative, unbiased prediction of ∆∆Gs with high throughput.

Here, we describe ThermoNet, a computational framework based on deep 3D convolutional neural networks (3D-CNNs) for predicting ∆∆Gs upon single-point mutation. We model the structure of each mutation assuming that single-point mutations introduce negligible perturbation to the overall architecture of protein structure. We treat protein structures as if they were 3D images with voxels parameterized using atom biophysical properties (27, 28). We leverage the power of the architecture of CNNs in detecting spatially proximate features. These local biochemical interaction detectors are then hierarchically composed into more intricate features with the potential to describe the complex and nonlinear phenomenon of molecular interaction. We address the bias in many previous methods towards predicting destabilization by training ThermoNet on a balanced data set generated through anti-symmetry-based data augmentation, i.e. for each mutation, we consider both the direct and reverse versions. We further demonstrate and address an unappreciated source of bias in previous performance estimates due to homology between training and evaluation sets. We show that ThermoNet achieves state-of-the-art performance comparable to previously developed methods on a widely used test set with minimal prediction bias. We also demonstrate the applicability of ThermoNet by showing that ThermoNet accurately predicts the ∆∆Gs of the missense mutations in two biologically important proteins, the p53 tumor suppressor protein and myoglobin and that ThermoNet-predicted ∆∆Gs of ClinVar missense variants fall within the experimentally observed range and are consistent with the expectations of a biophysical model of protein evolution.

## Results

### An overview of ThermoNet

To predict the ∆∆G of a point mutation, we take advantage of recent advances in deep learning for computer vision (29) and the successes of deep convolutional neural networks in biophysical problems (27, 28, 30–32). We treat protein structures as if they were 3D images with voxels parameterized using atom biophysical properties (27, 28) (Fig 1A,1B, and Table 1). For a given single-point mutation, ThermoNet requires that a 3D structure (either experimentally determined or modeled via homology modeling) of one of the alleles is available. As a first step, ThermoNet constructs a structural model for the mutant from the structure of the wild type using the Rosetta macromolecular modeling suite (Methods) (9, 33). ThermoNet assumes that the ∆∆G of a point mutation can be sufficiently captured by modeling the 3D physicochemical environment around the mutation site. It thus extracts predictive features by treating protein structures as if they were 3D images and voxelizing the space around the mutation site of both the wild-type structure and the corresponding mutant structural model (Fig 1C). Each voxel is parameterized with seven predefined rules (Table 1) to characterize the physicochemical nature of its neighboring atoms. The feature maps are then stacked to create a tensor with size [16,16,16,14] as input to the trained ensemble of ten deep 3D-CNNs, which generate a prediction of the ∆∆G the given mutation causes to the wild-type structure (Fig 1D, 1E, and Methods). Each of the component 3D-CNN models consists of three 3D convolutional layers with 16, 24, and 32 neurons respectively and one densely connected layer of 24 neurons (Fig 1E). These architectural hyperparameters (i.e. number of neurons in convolutional and densely connected layers and sizes of the input voxel grid) were tuned via five-fold cross-validation (Methods, Fig S1).

**Table 1.** Chemical property channels (AutoDock4 atom types) of a protein structure voxel

| Property | Rule |
| --- | --- |
| Hydrophobic | Aliphatic or aromatic carbon atoms |
| Aromatic | Aromatic carbon atoms |
| Hydrogen bond donor | Nitrogen, oxygen, sulfur atoms with lone-pair electrons |
| Hydrogen bond acceptor | Polar hydrogen atoms |
| Positive ionizable | Atoms with positive charge |
| Negative ionizable | Atoms with negative charge |
| Occupancy | All atom types |

**Fig 1.**
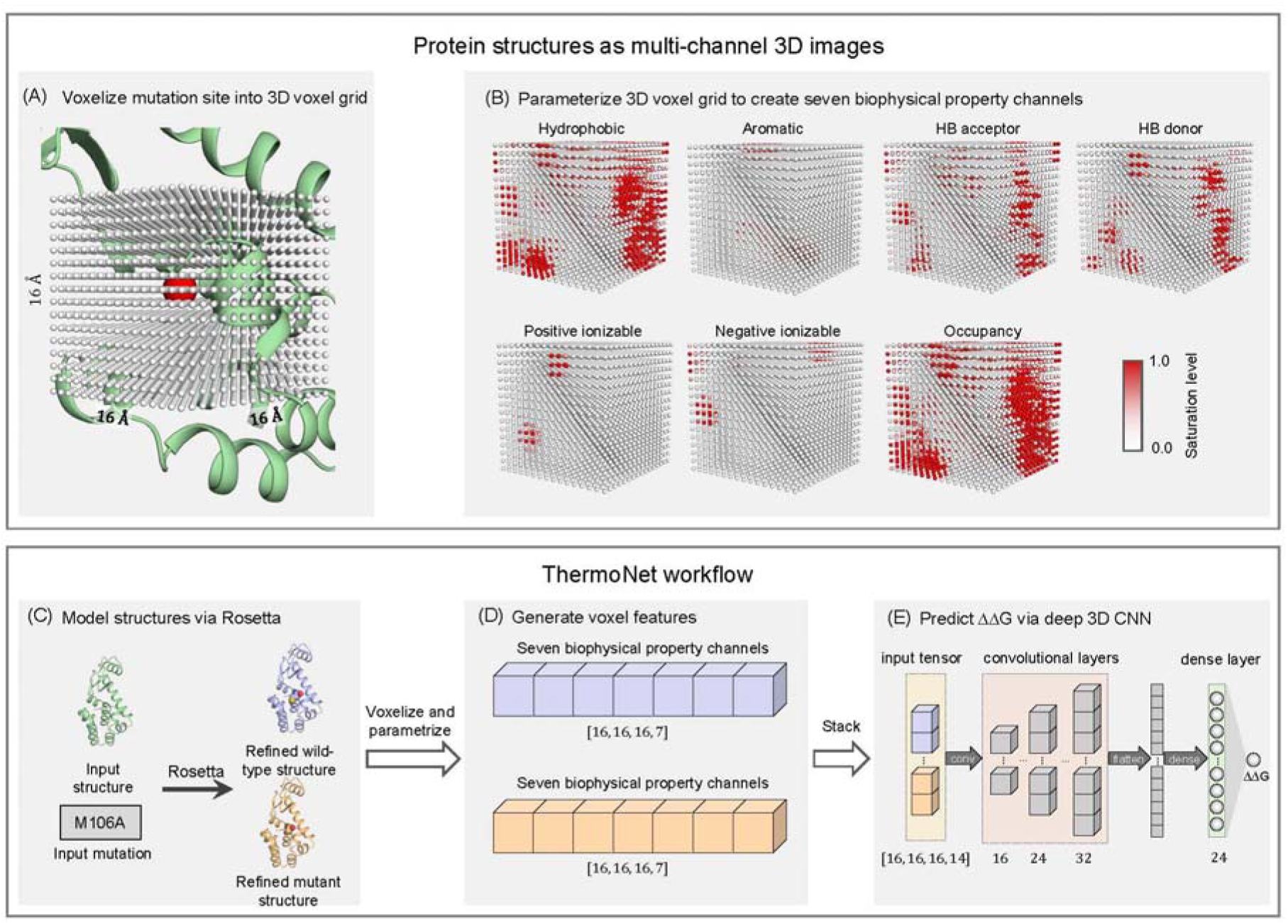
An overview of the ThermoNet computational framework. (A) Protein structures are treated as if they were 3D images. A 16 Å x 16 Å x 16 Å cubic neighborhood centered at the C_β_ atom (red sphere) of the mutated residue (or C_α_ atom in the case of a glycine) of an example protein (PDB ID: 1L63) is discretized into a 3D voxel grid at a resolution of 1 Å. Each voxel is represented by a gray dot. (B) Just as an RGB image has three color channels, the 3D voxel grid is parameterized with seven chemical property channels: hydrophobic, aromatic, hydrogen bonding donor, hydrogen bond acceptor, positive ionizable, negative ionizable, and occupancy. The saturation level of each voxel ranges from 0.0 to 1.0 and is colored accordingly (Methods). (C) To predict the change in thermodynamic stability caused by a given single-point mutation, ThermoNet calls Rosetta to refine the wild-type structure and to create a structural model of the mutant protein. (D) ThermoNet voxelizes the space around the mutation site of both the Rosetta-refined wild-type structure and the corresponding mutant structural model. Both the 3D voxel grid of the wild-type structure and that of the mutant model are parameterized accordingly to create two [16, 16, 16, 7] feature maps. (E) The feature maps are then stacked to create a [16, 16, 16, 14] tensor as an input to the trained deep 3D convolutional neural network. The final output of the network is the predicted ∆∆G the given mutation causes to the wild-type protein structure.

### Creating data sets for robust training and testing of ThermoNet

The ability of a machine-learning model to generalize can be overestimated when there is data leakage between the training set and the test set. In structural bioinformatics problems such as ∆∆G prediction, such data leakage can result when the training set contains proteins that are homologous to proteins in the test set. This is because the effects of different mutations in the same protein or homologous proteins are not necessarily independent. In most previous methods for ∆∆G prediction, this data leakage issue was not fully appreciated, and homologous proteins were present between training and test sets. For example, a recent method used randomly selected subsets of mutants from the widely used S2648 data set for training and testing (34). Not surprisingly, the training and test sets of shared 61 identical proteins (Supporting Information Table S1).

The issue of having mutations from the same protein in both training and test sets was addressed in developing the mCSM and INPS methods, where the cross-validation procedure ensured that mutations of the same protein remained together in either the training or test set (16, 19). While this was a step in the right direction, grouping mutations at the protein level is not sufficient to remove the homology between training and test sets. For example, we found substantial homology between 132 proteins within S2648 (Supporting Information Fig S2A). Thus, splitting the S2648 data set for training and testing at the protein level is likely to end up with shared homology between the splits. In fact, using the PISCES server (35) to remove redundancy from the S2648 data set resulted in only 104 non-redundant proteins (out of 132) at the level of < 25% sequence identity (Supporting Information Table S2). Homology is common among data sets used in training ∆∆G predictors; for example, many proteins in the VariBench (36) data set also share substantial homology (Fig S2B).

We further highlight that the widely used S2648 data set includes fourteen proteins from S^sym^ data set, which was previously used to evaluate a wide range of ∆∆G predictors (24, 37), and another eight proteins that are putative homologs to proteins in S^sym^ (Fig 2A, Supporting Information Table S3). In addition, a similar level of overlap also exists between the VariBench and Q3421 data sets and S^sym^ (Fig 2A Supporting Information Table S4 and S5). Thus, performance estimates on S^sym^ of methods trained using S2648 or parameterized using VariBench are likely to be overly optimistic. For example, in a recent publication, a version of INPS trained using a data set obtained by removing all proteins with > 25% sequence identity to proteins in S^sym^ showed substantially reduced performance (38).

**Fig 2.**
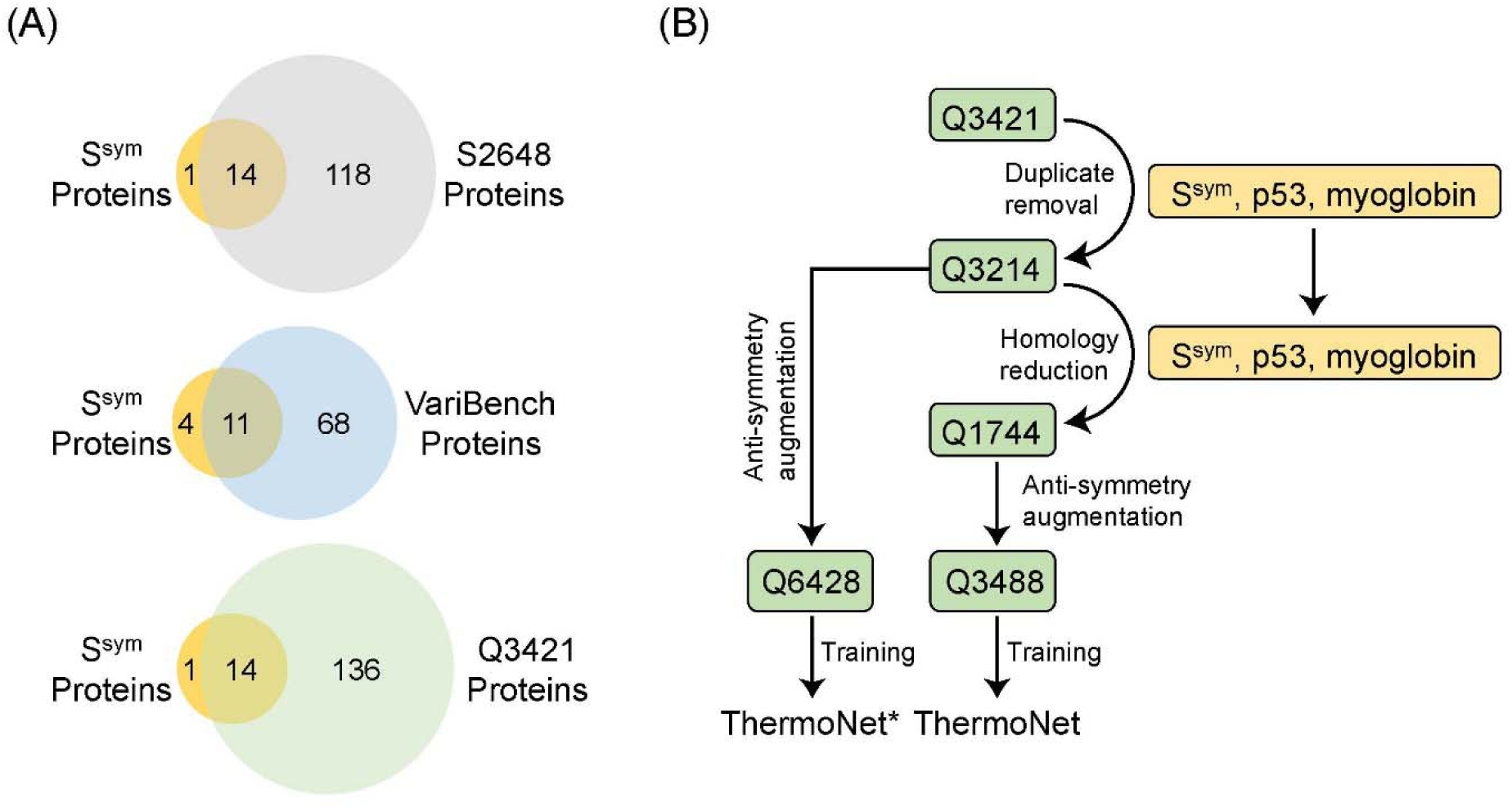
Data set curation and identification of shared homology. (A) Venn diagrams showing the amount of overlap at the protein level between three widely used training sets S2648, VariBench, and Q3421 for ∆∆G predictors and the S^sym^ test set. Numbers in these diagrams indicate protein counts. Upper panel and lower panel indicate that both S2648 and Q3421 share 14 identical proteins with S^sym^; middle panel indicates that VariBench and S^sym^ share 11 identical proteins. All three data sets share additional homology with S^sym^, which is presented in Supporting Information Table S3, S4, and S5, respectively. (B) Creating data sets for robust training and testing of ThermoNet. We started with the Q3421 set of 3421 mutations from 150 proteins. (Numbers in data set names indicate the number of unique mutations the data set contains.) After homology reduction and anti-symmetry data augmentation (Methods), this data curation workflow gives a training set of 3488 mutations with an equal representation of stabilizing and destabilizing changes and reduced homology to the S^sym^ test set. A separate data set called Q6428 was also created by augmenting the Q3214 data set before homology reduction to train ThermoNet*.

Thus, to train and evaluate ThermoNet, we implemented a rigorous procedure to reduce the sequence similarity between the training and test sets (S^sym^, p53, myoglobin) by removing duplicate data points and pruning protein-level homology (Fig 2B). Our rigorous pruning of the starting Q3421 data set (15) resulted in a data set consisting of 1,744 distinct mutations. This data set was then augmented by creating a reverse mutation data point for each of the 1,744 direct mutations according to the anti-symmetry property of ∆∆G (Methods), thus giving to a total of 3,488 data points for the training of ThermoNet (Fig 2B). While the pruning nearly halved the size of available training data, as discussed in the following section, models trained using the resulting augmented Q3488 data set will be less likely to have an overestimated performance when evaluated on S^sym^. Finally, to explore the influence of homology between training and test sets on estimates of model performance, we also augmented Q3214, the intermediate data set before the step of homology reduction, to train a different version of ThermoNet, called ThermoNet* (Fig 2B).

### ThermoNet achieves state-of-the-art performance on blind test set

We systematically compared ThermoNet and ThermoNet* with seventeen ∆∆G predictors on the S^sym^ balanced data set to evaluate their performance and degree of bias with respect to the ∆∆G anti-symmetry between direct and reverse mutations (Methods). A brief summary of the characteristics of these ∆∆G predictors and their references are given in Supporting Information Table S6. In short, the predictors are based on diverse features and strategies with some, like ThermoNet based only on structural information, while others like DDGun3D (37) and STRUM (15), integrate structural information with sequence and evolutionary features. The S^sym^ data set, which was constructed previously for assessing the biases of ∆∆G predictors (24). It consists of experimentally measured ∆∆G values for 342 direct and the corresponding reverse mutations (a total of 684 mutations) from fifteen protein chains for which the structures of both the wild-type and mutant proteins have been resolved by X-ray crystallography with a resolution of 2.5 Å or better. This data set is by construction balanced with respect to stabilizing and destabilizing mutations, thus enabling the evaluation of prediction bias. However, as noted in the previous section, many proteins in S^sym^ overlap or are homologous to proteins in commonly used training sets (Fig 2A).

To evaluate performance, we computed the root mean square error σ and the Pearson correlation coefficient *r* separately for direct and reverse mutations. We measured prediction bias by two statistics, the Pearson correlation coefficient *r*_*dir-rev*_ between the predictions for direct and those for reverse mutations and the δ value, defined as: *δ* = ∆∆*G*_*rev*_ + ∆Δ*G*_*dir*_. A perfectly unbiased predictor would give *r*_*dir-rev*_ = −1 and 〈*δ*〉 = 0 *kcal/mol*.

ThermoNet achieves strong prediction accuracy that is comparable for direct mutations (*r*_*dir*_ = 0.47 and *σ*_*dir*_ = 1.56 *kcal/mol*; Table 2, Fig 3A) and the corresponding set of reverse mutations (*r*_*rev*_ = 0.47 and *σ*_*rev*_ = 1.55 *kcal/mol*, Fig 3C). This suggests that ThermoNet did not overfit to direct mutations. The fractions of mutations for which the prediction error is within 0.5 *kcal/mol* and 1.0 *kcal/mol* are 36.3% and 58.8% for direct mutations and 36.5% and 58.5% for reverse mutations (Fig 3B and 3D). ThermoNet successfully reduces prediction bias with a near-perfect *r*_*dir-rev*_ (−0.96) and a negligible 〈*δ*〉 (−0.01) (Fig 3E). We also report the distribution of *δ*, since 〈*δ*〉 cannot distinguish large, but symmetric, bias from low bias (Methods). As shown in Fig 3F, 40.9% and 96.2% of mutations have a prediction bias < 0.1 *kcal/mol* and < 0.5 *kcal / mol*, respectively.

**Table 2.** Comparative analysis using the balanced blind test set S^sym^.

| Method | $\sigma_{\text{dir}}$ | $r_{\text{dir}}$ | $\sigma_{\text{rev}}$ | $r_{\text{rev}}$ | $r_{\text{dir-rev}}$ | $\langle\delta\rangle$ |
| --- | --- | --- | --- | --- | --- | --- |
| ThermoNet* | 1.42 | 0.58 | 1.38 | 0.59 | -0.95 | -0.05 |
| DDGun3D | 1.42 | 0.56 | 1.46 | 0.53 | -0.99 | -0.02 |
| DDGun | 1.47 | 0.48 | 1.50 | 0.48 | -0.99 | -0.01 |
| ThermoNet | 1.56 | 0.47 | 1.55 | 0.47 | -0.96 | -0.01 |
| PoPMuSiC<sup>sym</sup> | 1.58 | 0.48 | 1.62 | 0.48 | -0.77 | 0.03 |
| MAESTRO | 1.36 | 0.52 | 2.09 | 0.32 | -0.34 | -0.58 |
| FoldX | 1.56 | 0.63 | 2.13 | 0.39 | -0.38 | -0.47 |
| PoPMuSiC 2.1 | 1.21 | 0.63 | 2.18 | 0.25 | -0.29 | -0.71 |
| SDM | 1.74 | 0.51 | 2.28 | 0.32 | -0.75 | -0.32 |
| iSTABLE | 1.10 | 0.72 | 2.28 | -0.08 | -0.05 | -0.60 |
| I-Mutant 3.0 | 1.23 | 0.62 | 2.32 | -0.04 | 0.02 | -0.68 |
| NeEMO | 1.08 | 0.72 | 2.35 | 0.02 | 0.09 | -0.60 |
| DUET | 1.20 | 0.63 | 2.38 | 0.13 | -0.21 | -0.84 |
| mCSM | 1.23 | 0.61 | 2.43 | 0.14 | -0.26 | -0.91 |
| MUPRO | 0.94 | 0.79 | 2.51 | 0.07 | -0.02 | -0.97 |
| STRUM | 1.05 | 0.75 | 2.51 | -0.15 | 0.34 | -0.87 |
| Rosetta | 2.31 | 0.69 | 2.61 | 0.43 | -0.41 | -0.69 |
| AUTOMUTE | 1.07 | 0.73 | 2.61 | -0.01 | -0.06 | -0.99 |
| CUPSAT | 1.71 | 0.39 | 2.88 | 0.05 | -0.54 | -0.72 |

**Fig 3.**
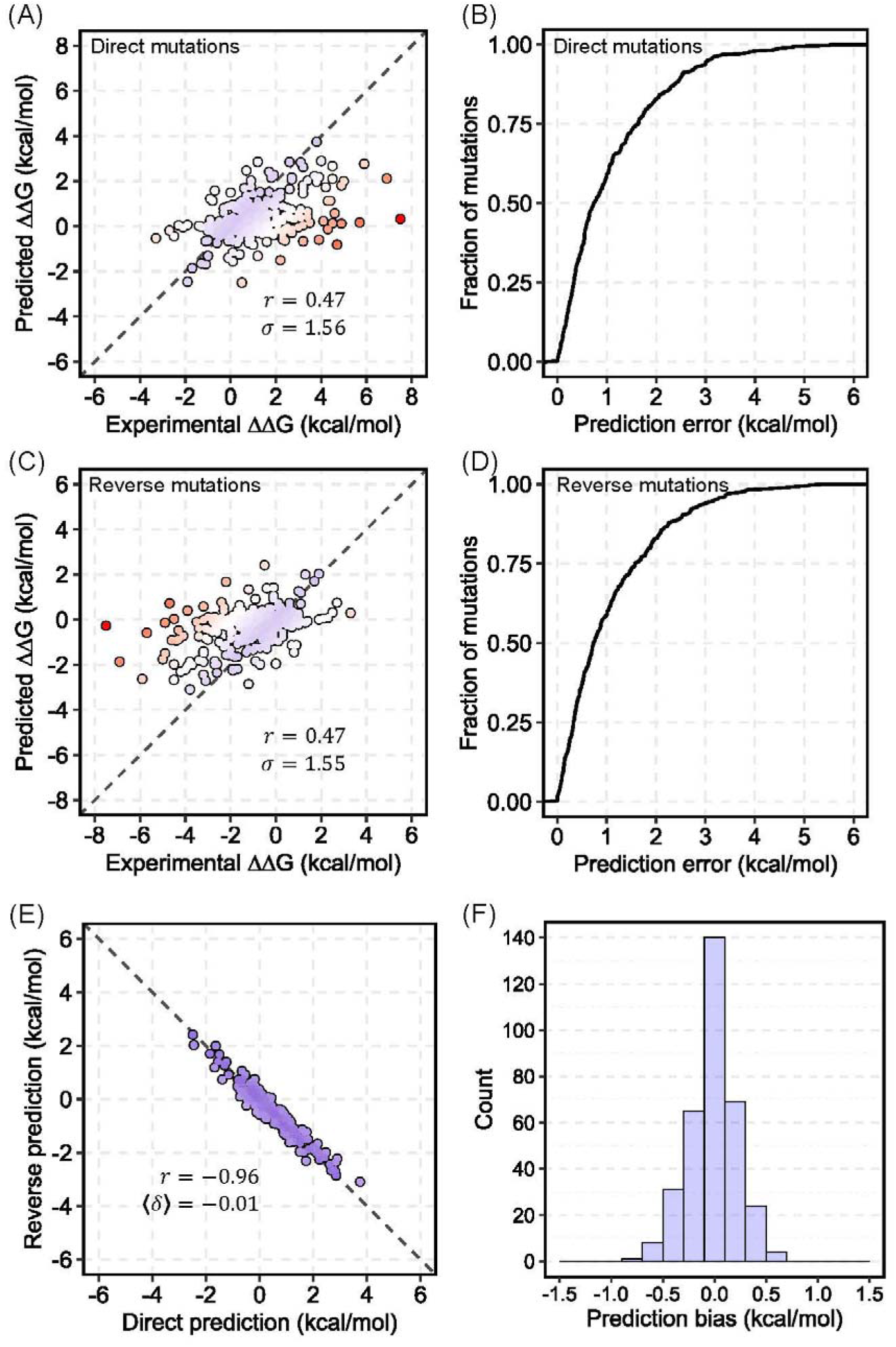
Performance of ThermoNet on the blind test set. (A) Performance of ThermoNet on predicting ∆∆G for direct mutations; The Pearson correlation coefficient (*r*) between predicted values and experimentally determined values is 0.47, and the root-mean-square deviation (*σ*) of predicted values from experimentally determined values is 1.56 kcal/mol. The dots are colored in gradient from blue to red such that blue represents the most accurate prediction and red indicates the least accurate prediction. (B) Cumulative distribution of ThermoNet prediction error on direct mutations. (C) Performance of ThermoNet on predicting ∆∆G for the reverse mutations (*r* = 0.47, *σ* = 1.55 kcal/mol). (D) Cumulative distribution of ThermoNet prediction error on reverse mutations. (E) Direct versus reverse ∆∆G values of all the mutations in the blind test set predicted by ThermoNet. A perfectly unbiased predictor would give *r* = −1 and 〈*δ*〉 = 0 *kcal/mol*. ThermoNet successfully reduces prediction bias with *r* = −0.96 and 〈*δ*〉 = −0.01 *kcal/mol*. (F) Distribution of ThermoNet prediction bias.

We also report the performance of ThermoNet*, different version of ThermoNet trained using the Q6428 data set augmented from the intermediate data set Q3214 before the step of homology reduction in our data set curation procedure (Fig 2B and Methods). ThermoNet* was trained in the exact same way as ThermoNet except that the homology of its training set Q3214 to S^sym^ was retained. Thus, the parameterization of ThermoNet* is comparable to previous methods that did not consider homology. As expected, evaluation of ThermoNet* on S^sym^ shows even better performance in *σ*_*rev*_ (1.38 vs. 1.55 kcal/mol) and *r*_*rev*_ (0.59 vs. 0.48) than ThermoNet, which was trained using the data set obtained after homology reduction (Table 2, Fig 2B, and Methods). Thus, the performance of many previously developed methods is likely to be substantially lower if they had been trained using a data set that shared no homology with S^sym^. In contrast, ThermoNet’s strong performance, even after removing homology reduction, suggests robust generalization in real-life applications.

Compared to other ∆∆G predictors, ThermoNet* achieves the best performance on reverse mutations, and the methods that outperform it on direct mutations all have substantial bias against reverse mutations (*σ*_*rev*_ > 2.09 kcal/mol and 〈 *δ* 〉 < −0.58). The seemingly good performance of many machine learning-based methods on direct mutations, but poor performance on reverse mutations suggests potential overfitting due to unbalanced training sets (24–26, 39). ThermoNet also performs well, but as a result of the reduction in performance due to removing homology between training and validation sets, the DDGun and DDGun3D methods outperform it on direct and reverse mutations. Unfortunately, it is not possible to retrain and evaluate all the other methods on the homology pruned training set, so we cannot directly compare the other methods to ThermoNet. Nonetheless, the fact that it still outperforms most suggests its utility and robustness.

### Structural models of reverse mutations are necessary for unbiased ∆∆G predictions

To evaluate whether the inclusion of the reverse mutations is necessary for the reduction in prediction bias, we trained a predictor following the same procedure for training ThermoNet but using a data set consisting of only the 1,744 direct mutations and their associated experimental ∆∆Gs (i.e. the Q1744 data set in Fig 2B). We applied this predictor to predict the ∆∆Gs of the direct and reverse mutations of the S^sym^ test set. As shown in Fig S3, ∆∆Gs of direct mutations predicted by these models correlate reasonably well (*r* = 0.47 and *σ* = 1.38 *kcal/mol*) with the experimental values and are comparable to the performance of the ensemble of networks trained using the balanced data set Q3488. In contrast, these models perform poorly (*r* = −0.06 and *σ* = 2.40 *kcal/mol*) in predicting the ∆∆Gs of the corresponding set of reverse mutations (Fig S3). This suggests that the models were biased toward the training set which is dominated by destabilizing mutations. This is confirmed by the strongly positive correlation between the predictions for direct mutations and those for reverse mutations and the large prediction bias (*r*_*dir-rev*_ = 0.35 and 〈*δ*〉 = 1.63 *kcal/mol*) (Fig S3). Compared to the performance of ThermoNet, which was trained using the balanced data set Q3488, the results highlight the necessity of a balanced data set for correcting prediction bias.

### Case studies: The p53 tumor suppressor protein and myoglobin

We further tested ThermoNet by predicting the ∆∆Gs of single-point mutations in the p53 tumor suppressor protein and myoglobin whose thermodynamic effects have previously been experimentally measured. The *TP53* tumor suppressor gene encodes the p53 transcription factor that is mutated in ~45% of all human cancers (Beroud and Soussi, 2003; Olivier et al. 2002). Unlike most tumor suppressor proteins that are inactivated by deletion or truncation mutations, single amino acid substitutions in p53 often modify DNA binding or disrupt the conformation and stability of p53 (Olivier et al., 2002). Myoglobin is a cytoplasmic globular protein that regulates cellular oxygen concentration in cardiac myocytes and oxidative skeletal muscle fibers by reversible binding of oxygen through its heme prosthetic group (40). The p53 data set consists of 42 mutations compiled in a previous study (16) within the DNA binding domain of p53. The myoglobin data set consists of 134 mutations scattered throughout the protein chain compiled in a previous study (41). We note that none of the mutations in these two data sets were present in the training set and that proteins that are likely to be homologous to p53 and myoglobin were also removed from the training set of ThermoNet (Fig 2B, Methods). We used published crystal structures of p53 (PDB ID: 2OCJ) and myoglobin (PDB ID: 1BZ6) to create one structural model for each of the mutations in these two data sets respectively using the *FastRelax* protocol in Rosetta (42). These predictions were compared directly with the experimentally determined thermodynamic effects (Fig 4).

**Fig 4.**
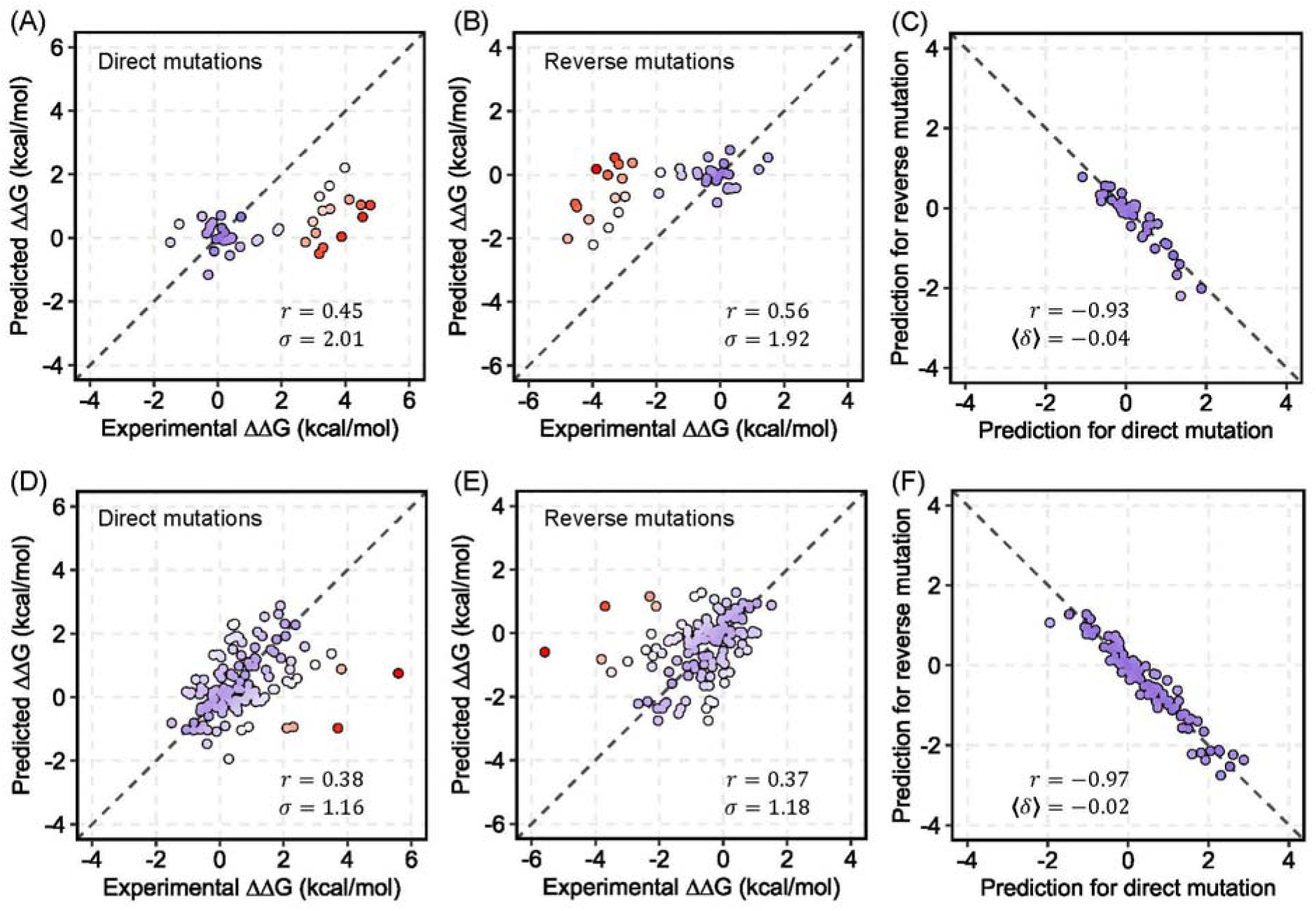
Predicted ThermoNet ΔΔG landscapes on p53 tumor suppressor protein and myoglobin. (A) Performance of ThermoNet on predicting ∆∆G for the direct mutations in p53 (*r* = 0.45, *σ* = 2.01 kcal/mol). (B) Performance of ThermoNet on predicting ΔΔG for the reverse mutations in p53 (*r* = 0.56, *σ* = 1.92 kcal/mol). (C) Direct versus reverse ∆∆G values of all p53 mutations predicted by ThermoNet (*r* = −0.93 and 〈*δ*〉 = −0.04 *kcal/mol*). (D) Performance of ThermoNet on predicting ∆∆G for the direct mutations in myoglobin (*r* = 0.38, *σ* = 1.16 kcal/mol). (E) Performance of ThermoNet on predicting ∆∆G for the reverse mutations in myoglobin (*r* = 0.37, *σ* = 1.18 kcal/mol). (F) Direct versus reverse ∆∆G values of all myoglobin mutations predicted by ThermoNet, with a Pearson correlation of *r* = −0.97 and 〈*δ*〉 = −0.02 *kcal/mol*. The dots are colored in gradient from blue to red such that blue represents the most accurate prediction and red indicates the least accurate prediction.

For p53, ∆∆Gs of both direct and reverse mutations predicted by ThermoNet correlate with the experimental measurements (*r* = 0.45 and 0.56) and have little bias (*r*_*dir-rev*_ = −0.93 and 〈δ〉 = −0.04 *kcal/mol*). However, the predicted ∆∆Gs of myoglobin mutations correlate less well (*r* = 0.38 and 0.37 for direct and reverse mutations respectively) compared to those of p53, though the bias is also low (*r*_*dir-rev*_ = −0.97 and 〈*δ*〉 = −0.02 *kcal/mol*). The poorer correlations for myoglobin are likely because the myoglobin data set consists of ∆∆G measurements obtained under various experimental conditions which ThermoNet does not explicitly account for. In fact, after excluding four data points (L29N, A130L, and two data points corresponding to A130K) with the biggest prediction error (> 3 kcal / mol), the Pearson correlations increase to 0.52 and 0.51 for direct and reverse mutations respectively. While the correlation between predicted and experimentally measured ∆∆Gs is not perfect, the predictions are generally conservative – no mutations with low measured ∆∆G are predicted to have a high ∆∆G. These results demonstrate the utility of ThermoNet as rapid unbiased predictor of ∆∆G for mutations in clinically relevant proteins.

We also compared ThermoNet with four other biophysics-based methods: FoldX, Rosetta, SDM, and CUPSAT. We were not able to include all methods because both p53 and myoglobin are already in the S2648 set that was used for training most of the other machine learning-based ∆∆G predictors and they are also in the VariBench (36) data set used to derive parameters of the DDGun model (37). Our comparison indicates that both FoldX and Rosetta predictions have a better correlation than ThermoNet while also reasonably anti-symmetric (Supporting Information Table S7 and S8). However, as shown in Table 2 and demonstrated in previous studies (24, 25), both FoldX and Rosetta are likely to show bias toward predicting destabilization when tested on larger data sets.

### ∆∆G landscape of ClinVar missense variants

Previous work has shown that variant deleteriousness can only be partially attributed to ∆∆G (43) and that both stabilization and destabilization can cause disease (44). We sought to use ThermoNet to obtain a less biased picture of the impact of benign and pathogenic variants on proteins stability. We applied ThermoNet to predict the ∆∆G distributions of pathogenic and benign missense variants in ClinVar, a widely used resource of medically important variants (45). For comparison, we also applied FoldX, a popular and freely available ∆∆G predictor (13, 22) to the ClinVar set. We first examined the overall predicted ∆∆G distribution of ClinVar variants. The ∆∆Gs of ClinVar variants predicted by ThermoNet range from −2.75 kcal/mol to +3.75 kca/mol (Fig 5A). Experimentally measured ∆∆Gs generally fall within −5 kcal/mol to +5 kcal/mol (13, 46); thus, ThermoNet’s predictions are consistent with the range of observed values. In contrast, the ∆∆Gs of the same set of variants predicted by FoldX range from −6.64 kcal/mol to +57.2 kcal/mol (Fig 5A), and 15.2% of ∆∆Gs predicted by FoldX are outside the expected range of −5 kcal/mol to +5 kcal/mol.

**Fig 5.**
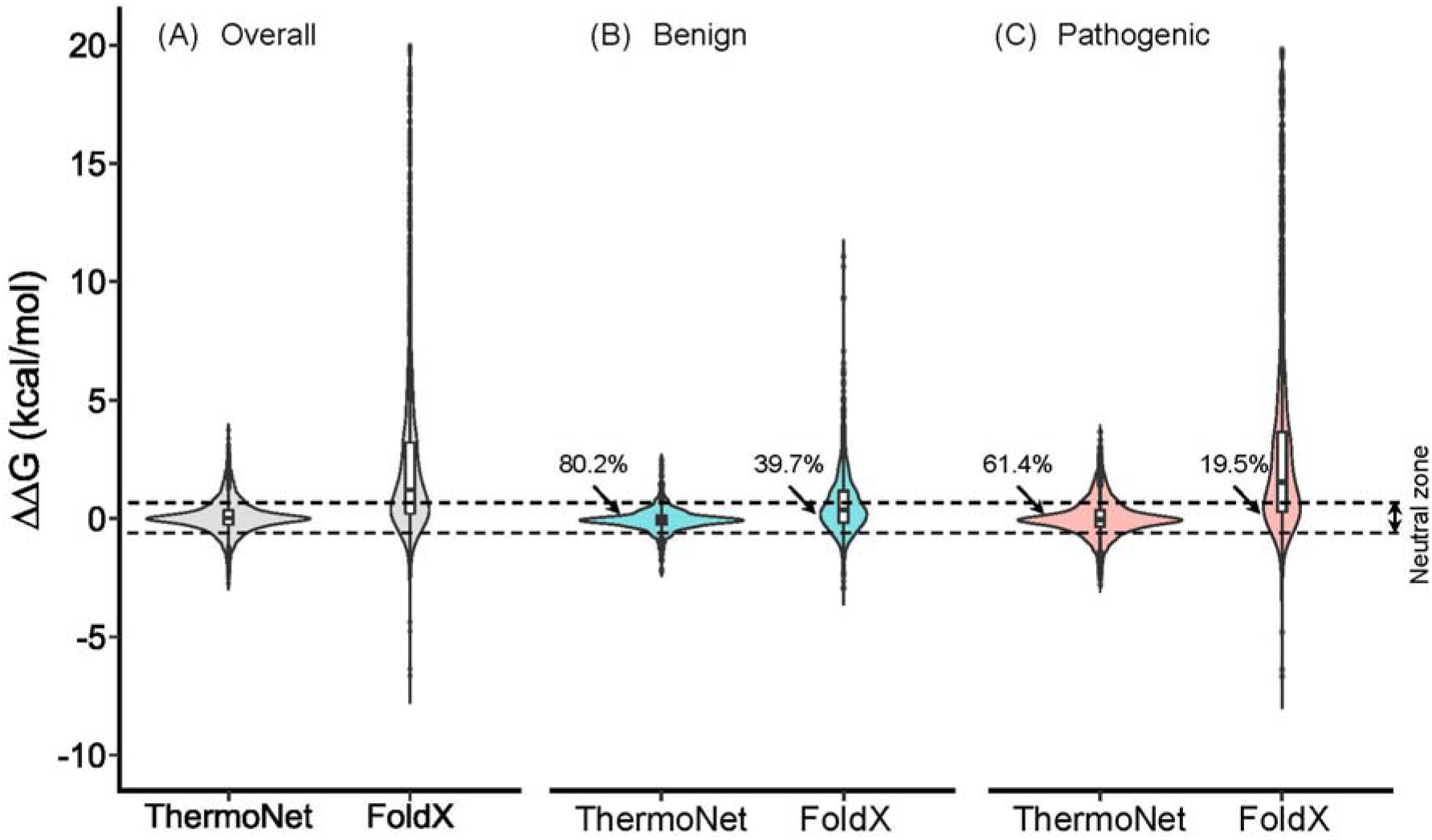
Predicted ∆∆G distributions of ClinVar missense variants. (A) The overall ∆∆G distributions of ClinVar variants predicted by ThermoNet and FoldX. ThermoNet’s predictions are consistent with the expected range based on experimentally determined ∆∆G values (−5 kcal/mol to +5 kcal/mol). In contrast, more than 15% of ∆∆Gs predicted by FoldX are outside the expected range. (B) The ∆∆G distributions for ClinVar benign variants predicted by ThermoNet and FoldX. (C) The ∆∆G distributions of ClinVar pathogenic variants predicted by ThermoNet and FoldX. The ∆∆Gs of 80.2% of benign variants predicted by ThermoNet fall within the neutral zone (−0.5 to +0.5 kcal/mol, region between dashed lines), in which variants are not expected to influence fitness. FoldX only predicted 39.7% of benign variants to be in the neutral zone. Further, the ∆∆Gs of pathogenic variants predicted by ThermoNet suggest pathogenic variants are nearly equally likely to be stabilizing (47.3%) as destabilizing (52.7%). In contrast, FoldX predicted that 83.2% of pathogenic variants are destabilizing. Variants for which FoldX ∆∆G is > 20 kcal/mol are omitted for clarity. Percentages represent the fractions of variants whose ∆∆Gs are predicted to be in the neutral zone.

∆∆Gs predicted by ThermoNet are also consistent with the expected ∆∆G distributions according to a biophysical model of protein evolution (46). This model hypothesizes that fitness is a non-monotonic, concave function of protein stability, meaning that fitness decreases with increasing deviation from an optimal stability. The model also suggests that there is a “neutral zone” of 1.0 kcal/mol around the optimal stability in stability space, and mutations whose impact on stability fall within the neutral zone will have little effect on fitness (46). We thus reasoned that the ∆∆Gs of benign variants should fall within a narrow range from −0.5 kcal/mol to +0.5 kcal/mol and that pathogenic variants should be equally likely to be destabilizing or stabilizing (46). To test this hypothesis, we examined the ∆∆G distributions of pathogenic and benign variants separately. The ∆∆Gs of 80.2% of benign variants predicted by ThermoNet fall within the neutral zone, whereas FoldX only predicted 39.7% of benign variants to be in the neutral zone (Fig 5B). Further, the ∆∆Gs of pathogenic variants predicted by ThermoNet suggest pathogenic variants are nearly equally likely to be destabilizing (52.7%) as they are to be stabilizing (47.3%) (Fig 5C). In contrast, FoldX predicted that 83.2% of pathogenic variants are destabilizing. As already demonstrated in previous studies, the bias is likely because FoldX was parameterized on an experimental ∆∆G data set dominated by destabilizing mutations (23–25).

ThermoNet’s predictions are also consistent with the fact that variant pathogenicity can only be partially attributed to impacts on protein stability. ThermoNet predicts that 61.4% of pathogenic variants that have ∆∆Gs within the neutral zone and 19.8% of benign variants have ∆∆Gs outside the neutral zone (Fig 5B and 5C). This is expected based on previous biochemical characterizations of pathogenic mutations. For example, Bromberg *et al.* collected 66 mutations with experimentally measured ∆∆G and functional annotations from the literature. The ∆∆Gs of this set of mutations range from −4.3 to 4.96 kcal/mol and the authors found that 31% of mutations affecting function had ∆∆Gs within the neutral zone while 19% functionally neutral mutations had ∆∆Gs outside the neutral zone (43).

## Discussion

Accurate modeling of protein thermodynamic stability is a complex task due to the delicate balance between the different thermodynamic state functions that contribute to protein stability (1). The primary goal of this paper is to present a novel application of deep 3D convolutional neural networks to a fundamental challenge in structural bioinformatics: predicting changes in thermodynamic stability upon point mutation. We formulated the problem of ∆∆G prediction from a computer vision perspective and took full advantage of the power of the constrained architecture of convolutional neural networks in detecting spatially proximate features (29). We developed ThermoNet, a method based on deep 3D convolutional neural networks, to predict ∆∆G upon point mutation. We showed that ∆∆G can be predicted from protein structure with reasonable accuracy using deep 3D convolutional neural networks without manual feature engineering. While ThermoNet achieved comparable performance to previous methods on direct mutations, it performed better on reverse mutations than most methods by a large margin, and remarkably, reduced the magnitude of prediction bias.

In addition to introducing ThermoNet, we also address two methodological challenges in the development and evaluation of computational methods for ∆∆G prediction: lack of anti-symmetry and data leaks due to homology between proteins in the training and evaluation sets. Previously, it was shown that the lack of anti-symmetry in ∆∆G prediction can be effectively addressed either by using input features that are anti-symmetric by construction (19, 37, 38, 47, 48) or by training the predictor using both direct and reverse mutations (19, 24). In addition, when the statistical model is parametric, one may identify the terms that are responsible for breaking the symmetry and make correction accordingly (48). However, when the predictor is nonparametric, meaning that the terms of the statistical learning model are not established a priori, the only way is to train the model with data set balanced with direct and reverse mutations so that it learns the anti-symmetry (24). For structure-based ∆∆G predictors, this approach requires knowing the 3D structures of both the wild type and mutant protein. While mutant structures determined via experimental techniques are scarce, in this work, we demonstrated that mutant protein structures obtained through molecular modeling-based data augmentation can also be effectively used as substitutes for experimental structures to remedy the lack of anti-symmetry in ∆∆G prediction.

Recently, the potential for data leak between training and testing due to the inclusion of mutations from the same proteins has been appreciated (16, 19); however, the effects of including mutations from homologous proteins in training and validation sets is less appreciated and understood. Our results suggest that that such homology can influence performance estimates. The ThermoNet* model, which was trained before homology reduction, achieved stronger performance than the ThermoNet model trained after homology reduction (Fig 2B). In real-world applications, there will be homology between proteins used to train prediction models and the proteins to which they are applied. However, given the relatively small number of proteins included in commonly used training sets and the fact that they are not representative of the full diversity of protein folds and functions, we believe that the inclusion of mutations from proteins with shared evolutionary histories is likely to bias performance estimates. In the future, it will be valuable to explore this issue further and construct training sets that reflect the evolutionary relationships expected in various applications.

ThermoNet treats protein structures as if they were 3D images, and it takes as input a 3D grid of voxels parameterized with seven biophysical property channels. As such, this approach bypasses the tedious processes of manual feature engineering and feature selection that, if not done correctly, can often lead to over-optimistic estimation of model performance. The locally constrained deep convolutional architecture likely allows the system to model the complex, non-linear nature of molecular interactions. Recently, a spherical convolutional architecture in which concentric voxel grids parameterized by atom masses and charges were used as input to predict the ∆∆Gs of direct mutations with good accuracy (49). Thus, together with the current work, these results demonstrate the potential of deep convolutional neural networks for predicting biophysically meaningful information from protein structures and holds promise for protein engineering.

The fact that our approach relies on the availability of experimental structures or homology models and the 3D nature of the convolutional neural network create two limitations. First, while protein structures are being determined at an unprecedented pace, the fraction of the human proteome with available experimental structure is estimated to be around 20% (50). Even when all the proteins whose structures can be modeled reliably are considered, only ~70% of the human proteome will have structural coverage (50). As the structures of many proteins can only be partially modeled, the space of the human proteome that one can apply ThermoNet to will be less than 70%. Second, compared to the 2D version with the same architecture, ThermoNet has four times more parameters: three convolutional layers of 16, 24, and 32 neurons respectively, and one dense layer with 24 neurons that takes an input tensor of the shape [16, 16, 16, 14] has 133,273 parameters. Training deep 3D CNNs is very demanding, requiring more GPU memory and more training data to avoid overfitting. While we demonstrated the potential of deep 3D CNNs in modeling ∆∆G of proteins, the relatively little training data available raises the question of whether deep 3D CNNs can model ∆∆Gs and related thermodynamic properties at experimental accuracy. Nonetheless, the increasing adoption of deep mutational scanning techniques for systematic study of the molecular effects of mutations is generating an unprecedented amount of data. Furthermore, given rapid increases in GPU power and the number of structures of proteins and their complexes determined, we expect that deep 3D CNNs will be successfully applied to provide solutions to many biophysical problems such as modeling the impact of mutation on protein-protein, protein-DNA as well as protein-RNA interactions.

## Methods

### Protein thermodynamic stability

The thermodynamic stability ∆G of the folded form of a two-state protein, which is its Gibbs free energy of folding, is defined in relation to the concentration of folded [*folded*] and the concentration of unfolded [*unfolded*] forms:

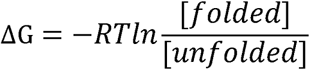

where T is the temperature, and R is the gas constant. More stable proteins, meaning that a higher fraction of the protein is in the folded form, have more negative values of ∆G. The impact of mutations on protein stability, ∆∆G, is defined in terms of the change in ∆G between the wild-type and mutant proteins:

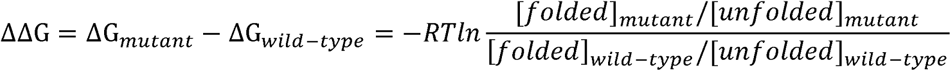

such that a destabilizing mutation has a positive ∆∆G, whereas a stabilizing mutation has a negative ∆∆G. The values of ∆∆G resulting from single-point mutations usually range from −5 to 5 kcal/mol (13). A mutation that destabilizes a typical protein (∆G = −5 kcal/mol) by 1 kcal/mol will reduce the equilibrium constant for the folding reaction of this protein by a factor of 5.1 at physiological temperature (310 K).

### Thermodynamics of direct and reverse mutations

Consider a pair of proteins whose sequences differ only at a single position where the amino acid is *X* in one protein and *Y* in the other. Let the Gibbs free energies of folding of this pair of Consider a pair of proteins whose sequences differ only at a single position where the amino proteins be ∆*G*_*X*_ and ∆*G*_*Y*_ respectively. For such a pair of proteins, one can think of the protein *Y* as being generated by a “direct” mutation at the sequence location from amino acid *X* to *Y* and the change in the Gibbs free energy of folding caused by *X* to *Y* mutation is ∆∆*G*_*X→Y*_ = ∆*G*_*Y*_ − ∆*G*_*X*_. One may also think of the protein *X* as being generated by a “reverse” mutation from amino acid *Y* to *X* and the change in the Gibbs free energy of folding caused by this reverse mutation is ∆∆*G*_*Y→X*_ = ∆*G*_*X*_ − ∆*G*_*Y*_ = −∆∆*G*_*X→Y*_. A well-performing, “self-consistent” method for predicting ∆∆*G*s would not only give accurate ∆∆*G* predictions for the direct mutations, but also previously developed ∆∆*G* predictors (23–25). for the reverse mutations. The self-consistency requirement has been largely ignored by previously developed ∆∆*G* predictors (23–25).

### Data sets and symmetry-based data augmentation

The data set used to train ThermoNet was derived from the Q3421 data set compiled in a previous study (15). The Q3421 data set contains 3,421 distinct single-point mutations in 150 proteins collected from the ProTherm database (51). The impact of these mutations on the stability of the protein structure have been measured experimentally and expressed quantitatively as ∆∆G values. We first excluded those mutations from the Q3421 data set that were also in the S^sym^ test set (see following). To reduce the sequence similarity between proteins in the training set of ThermoNet and the proteins it was tested on, we also removed all proteins that are likely homologous (BLAST evalue < 0.001) to p53, myoglobin, and proteins in S^sym^ from Q3421. Estimation of homology was accomplished by running the blastp program (52) using protein sequences in the S^sym^ data set and the sequences of p53 and myoglobin as queries against protein sequences in Q3421. Our rigorous pruning of the Q3421 data set resulted in a final data set consisting of 1,744 distinct mutations in 127 proteins. This data set was augmented by creating a reverse mutation data point for each of the 1,744 direct mutations, thus giving to a total of 3,488 data points for the training of ThermoNet. The data set was randomly divided into ten equally sized, mutually exclusive subsets each consisting of 10% of the direct mutations and the corresponding 10% of reverse mutations. In the training of each component model of ThermoNet, nine subsets were combined to form a training set and the remaining one subset was used as a validation set. The data set used to test ThermoNet and to compare it with fifteen previously developed ∆∆G predictors was a common, balanced data set called S^sym^ consisting of 342 pairs of proteins with known crystal structures (24). The members forming each pair differ at only a single position in the protein sequence. The ∆∆G values of the 342 direct mutations have been experimentally measured and the ∆∆G values of the corresponding 342 reverse mutations were assigned using anti-symmetry.

### Modeling mutant structures

We treat each mutation as a pair of proteins whose sequences differ only at a single sequence position. For each pair of proteins, we designate the one whose structure has been experimentally resolved as protein X and the other as protein Y. Structures of the X proteins were collected from the Protein Data Bank (53) and were relaxed in the Rosetta all-atom energy function ref2015 (54) using the Rosetta *FastRelax* protocol (42). To prevent large-scale conformational shift from the input PDB structure, atoms were constrained to their starting locations with a harmonic penalty potential during relaxation. The same Rosetta *FastRelax* protocol was also employed to create structural models for each of the Y proteins from the corresponding relaxed structure of the X protein by supplying a Rosetta `resfile` specifying the mutation *X* → *Y* to make. The structures of both proteins of each protein pair in the test set were collected from the Protein Data Bank (53) and were also relaxed using the same Rosetta *FastRelax* protocol.

### Voxelization of the neighborhood of mutation site

We treated each protein structure as a collection of volume elements (voxels) in 3D space. Just as a pixel element in an image has color channels, we parameterized a voxel in a protein structure by a set of *k* chemical property channels: [*ν*_1_, *ν*_1_, …, *ν*_*k*_] where the value *ν*_*i*_ of each property channel indicates the level of saturation of property *i* at this voxel (Fig 1A and 1B). For each mutation (a pair of proteins), we superimposed the mutant structure onto the wild-type structure such that the root-mean-squared distance between them is minimized and collected a grid of 16 × 16 × 16 voxels from both structures. We parameterized each voxel with seven property channels each of a distinct chemical nature according to AutoDock4 atom types (55) as in the work of Jimenez et al. (28) (Table 1). This resulted in a tensor of the shape [16,16,16,7] for a single structure and a tensor of the shape [16,16,16,14] when the two tensors from both structures are concatenated to represent the mutation. The grid was centered at the C_β_ atom of the mutation site amino acid (or C_α_ atom if it’s a glycine) where each voxel is a unit cube whose sides are 1 Å long. The level of saturation *f(d)* of each property channel at each voxel is determined by the van der Waals radius *r*_*vdw*_ of the atom designated to have that property and its distance *d* to the center of the voxel through the following formula:

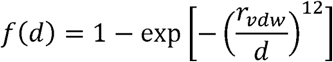

The computation of tensors from protein structures was performed using routines implemented in the HTMD Python library (version 1.17) for molecular simulations (56) and a Python program for creating a data set from a list of mutations is provided in Supplementary Code.

### Model architecture

Convolutional neural networks are a type of deep-learning model commonly used in computer vision applications. They have recently proven to perform well in residue contact prediction (57–60) and protein tertiary structure prediction (61–64). We selected CNNs for protein structure-based ∆∆G prediction because we formulated this problem as a computer vision problem by treating protein structures as if they were 3D images. Each of the component models of ThermoNet features a sequential organization of three 3D convolutional layers (Conv3D), one 3D max pooling layer (MaxPool3D), followed by one fully connected layer (Dense) (Fig S1). Convolutions operate over 4D tensors, called feature maps, with three spatial axes (length, width, and height) as well as a depth axis. The convolution operation extracts 3D patches of shape [3,3,3] with stride 1 from its input feature map and applies the same transformation to all patches, producing an output feature map. This output feature map is still a 4D tensor: it has a length, height, and a width whose values are determined by the shape of convolution patches and size of the stride, but its depth, which is also called number of filters, is a hyperparameter of the layer (see the section on hyperparameter search below). The number of filters in the three Conv3D layers were 16, 24, and 32 respectively. All convolution operation outputs from each Conv3D layer are transformed by the rectified linear activation function (ReLU). The transformed outputs from the last Conv3D layer are pooled by taking the maximum activation of each 2 × 2 × 2 grid. The max-pooled activations are then flattened into a 1D vector of features which are fully connected with a dense layer of 24 ReLU units. This model architecture was implemented using Keras (65) with TensorFlow (66) as the backend.

### Training ThermoNet

ThermoNet is an ensemble of ten deep 3D convolutional neural networks, each trained to perform the best on a validation set. Each of the component models was trained on nine subsets (collectively known as the training set), and its generalization performance was monitored on the remaining one subset (validation set). This process was iterated ten times each using a different one of the ten subsets as the validation set and the remaining nine subsets as the training set. We employed this procedure to obtain an ensemble of ten models as model ensembling has been suggested to produce better predictions (67). Evaluation of this ensemble was performed on a separate test set (see below). All component models of ThermoNet were trained using the Adam optimizer (68) for 200 epochs with default hyperparameters (maximum learning rate = 0.001, *β*_1_ = 0.9, *β*_2_ = 0.999). Kernel weights of the model were initialized using the Glorot uniform initializer and updated after processing each batch of eight training examples. The mean squared error (MSE) of the predicted ∆∆G values from the experimental measurements was used as the loss function during training. When training a deep neural network, one often cannot predict how many epochs will be needed to get to an optimal validation loss. We monitored the MSE of the predictions on a separate validation set consisting of 10%variants randomly selected from the training set during training. Training was stopped when the MSE on the validation set stopped decreasing for ten consecutive epochs. To regularize the model, the dense layer was placed between two dropout layers with a dropout rate of 0.5 in each layer (Fig S1). Each of the final component models was the one that produced the lowest MSE on the validation set. The final predicted ∆∆G value is the average of predictions from the ten models.

### Hyperparameter search

The design of deep CNNs entails many architectural choices to account for number of hidden convolutional layers and fully connected layers, number of filters, filter size, strides, padding, dropout rate among many other hyperparameters. We initially created a voxel grid with size 16 × 16 × 16 at a resolution of 1 Å for each chemical property channel following the procedure described in (27) to cross-validate our network architectures. Considering the limited training data set available in our study, we tried some smaller architectures with cross-validation to decide the optimal one, rather than simply adapting the widely used, much larger, network architectures in computer vision applications. We restricted our deep CNNs to have three convolutional layers and one fully connected while considering several hidden layer sizes. Our results from five-fold cross-validation suggest that the architecture of the 16 × 24 × 32 convolutional configuration combined with a fully connected layer of size 24 achieved the best performance (Fig S1). An additional consideration for our deep 3D CNN architecture is the dimension of the local box. The size of the local box specifies the structural information accessible by the network and therefore is a hyperparameter of our method. We cross-validated the voxelization scheme with grid of sizes 8 × 8 × 8, 12 × 12 × 12, 16 × 16 × 16, and 20 × 20 × 20 at a resolution of 1 Å. All voxelization schemes draw a cubic box around the mutation site with lateral length of *l* Å where *l* equals 8, 12, 16, or 20 Å. Our results from five-fold cross-validation indicate that the voxel grid with size 16 × 16 × 16 gives the best prediction performance (Fig S1).

### Performance evaluation

The following measures were adopted to evaluate the performance of ThermoNet and to facilitate comparison with previously developed methods. The primary measures for evaluating prediction accuracy were the Pearson correlation coefficient (*r*) between experimental and predicted ∆∆*G*s and the root-mean-squared error (*σ*) of predictions. For a set of *n* data points (*x*_*i*_, *y*_*i*_), the formula for calculating *r* and *σ* are defined as follows:

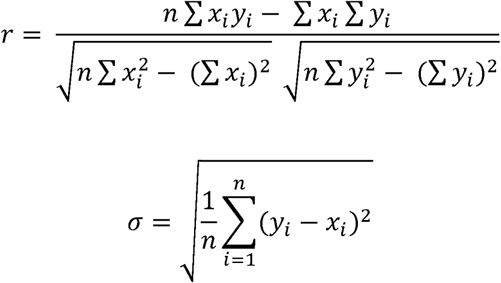

where the (*x*_*i*_, *y*_*i*_) tuple denotes the experimental and predicted ∆∆*G* values of mutation *i*, respectively, and *n* denotes the number of mutations in the data set. The measures for evaluating prediction bias were the Pearson correlation coefficient between the predictions for direct mutations and those for reverse mutations and the parameter which is defined as: *δ* = ∆Δ*G*_*inv*_ + ∆Δ*G*_*dir*_ and was previously used to quantify prediction bias (23). An unbiased predictor should have *δ* = 0 for every mutation. The average of *δ*, 〈*δ*〉, taken over all mutations in the S^sym^ data set was used in two previously studies to quantify prediction bias (24, 37). While we report 〈*δ*〉 in this work, we note that 〈*δ*〉 is flawed because biases toward opposite directions will be washed out when summed. To give a more transparent presentation of prediction bias, we also plot the distribution of *δ* and report the average of absolute bias, i.e. 〈|*δ*|〉.

### ClinVar variants

We retrieved the ClinVar database (45) in VCF format on August 15, 2019 and ran the VCF file through the Variant Effect Predictor (version 97) (69) to annotate the consequences of all ClinVar variants. We created a set of missense variants that can be mapped to protein structures to demonstrate the applicability of ThermoNet to clinically relevant variants. Our evaluation set consists of solely ClinVar missense variants that are labeled as “pathogenic” or “likely pathogenic” for true positive (pathogenic) variants and “benign” or “likely benign” for true negative (benign) variants. All variants are required to have a review status of at least one star and no conflicting interpretation. Any ClinVar variant designated as “no assertion criteria provided”, “no assertion provided”, “no interpretation for the single variant”, or not covered by either a structure or homology model was excluded from the evaluation set. Due to the dependency of ThermoNet on 3D structures, we also require variants in the evaluation set to be mappable to available protein structures. The residue-level mapping of ClinVar variants onto protein structures was based on the SIFTS resource that provides residue-level mapping between UniProt and Protein Data Bank (PDB) entries (70). Collectively, these restrictions resulted in 3,510 pathogenic variants and 950 benign variants that can be mapped to experimental structures deposited in the PDB (53). The mapped variants along with PDB IDs can be found in our GitHub repository at https://github.com/gersteinlab/ThermoNet.

## Author contribution

B.L. and M.B.G. conceived the study. B.L. implemented ThermoNet. B.L. and Y.T.Y performed the analysis. B.L. performed the work on predicting the ∆∆G distributions of ClinVar variants using ThermoNet at Vanderbilt University with the help from J.A.C. M.B.G. and J.A.C. supervised the project. B.L. wrote the manuscript with help from all authors.

## Code and data availability

ThermoNet source code and raw data supporting the analysis of this work is available at https://github.com/gersteinlab/ThermoNet.

## Acknowledgement

This work was supported by NSF award DBI1660648 (M.B.G.), NIH awards R35 GM127087 (J.A.C.) and R01 GM126249 (J.A.C.), and an American Heart Association Postdoctoral Fellowship 20POST35220002 (B.L.).The authors would also like to thank the Center for Research Computing at Yale University and the Advanced Computing Center for Research and Education at Vanderbilt University for supporting high-performance computing.

