## Supporting Information for "Predicting changes in protein thermodynamic stability upon point mutation with deep 3D convolutional neural networks"

Table S1. PDB IDs of Identical proteins between PoPMuSiC-2.0 training and test sets.

| 1aj3A | 1cunA | 1hmkA | 1mjcA | 1titA | 2nvhA |
| --- | --- | --- | --- | --- | --- |
| 1akyA | 1dktA | 1hmsA | 1msiA | 1ttqA | 2rn2A |
| 1aonU | 1e65A | 1ifcA | 1oiaA | 1uzcA | 2trxA |
| 1apsA | 1ey0A | 1igvA | 1p2pA | 1yyjA | 3glyA |
| 1bniA | 1fnaA | 1ihbA | 1qlpA | 1zg4A | 3mbpA |
| 1btaA | 1ftgA | 1imqA | 1rg8A | 1znjB | 3pgkA |
| 1bvcA | 1g4iA | 1k9qA | 1risA | 2a36A | 3silA |
| 1c9oA | 1h7mA | 1lniA | 1rn1C | 2driA | 4lyzA |
| 1ceyA | 1hfzA | 1lz1A | 1rtbA | 2immA | 5dfrA |
| 1cseI | 1hmeA | 1mgrA | 1shfA | 2lzmA | 5ptiA |
| 1cspA |  |  |  |  |  |

Table S2. PDB IDs of non-redundant proteins returned by submitting proteins in the S2648 data set to the PISCES server.

| 1a43A | 1dktA | 1jiwI | 1qlpA | 1yyjA |
| --- | --- | --- | --- | --- |
| 1aepA | 1e65A | 1k9qA | 1qm4A | 1zg4A |
| 1ag2A | 1ey0A | 1kcqA | 1qndA | 2a36A |
| 1akyA | 1fkjA | 1kdxA | 1rg8A | 2abdA |
| 1am7A | 1fnaA | 1ke4A | 1rhgA | 2driA |
| 1amqA | 1ftgA | 1kfwA | 1risA | 2lzmA |
| 1aonU | 1fvkA | 1lbiA | 1rn1C | 2nvhA |
| 1apsA | 1g4iA | 1lniA | 1ropA | 2ocjA |
| 1arrA | 1g6nA | 1lucA | 1sakA | 2rn2A |
| 1azpA | 1h7mA | 1lveA | 1shgA | 2trtA |
| 1b26A | 1hk0X | 1lz1A | 1supA | 2trxA |
| 1b8eA | 1hmeA | 1mbgA | 1tenA | 2ts1A |
| 1boyA | 1htiA | 1msiA | 1titA | 3ecaA |
| 1btaA | 1huuA | 1n0jA | 1tpkA | 3glyA |
| 1bvcA | 1ietA | 1oh0A | 1ttqA | 3hhrA |
| 1c9oA | 1ifcA | 1oiaA | 1tyvA | 3mbpA |
| 1cahA | 1igvA | 1oncA | 1ubqA | 3silA |
| 1ceyA | 1ihbA | 1pdoA | 1uzcA | 3ssiA |
| 1chkA | 1imqA | 1pgaA | 1vqbA | 5croA |
| 1cseI | 1io2A | 1pohA | 1witA | 5dfrA |
| 1cunA | 1iroA | 1qgvA | 1yu5X |  |

Table S3. Proteins in the S2648 data set (Subject) that are either identical or likely to be homologous to proteins in the S^sym^ data set (Query).

| Query | Subject | %ID | Length | # Mismatches | Query start | Query end | Subject start | Subject end | E-value |
| --- | --- | --- | --- | --- | --- | --- | --- | --- | --- |
| 1amqA | 1amqA | 100 | 396 | 0 | 1 | 396 | 1 | 396 | 0 |
| 1bniA | 1bniA | 100 | 108 | 0 | 1 | 108 | 1 | 108 | 1.97E-80 |
| 1bniA | 1mgrA | 37.3 | 59 | 35 | 51 | 107 | 35 | 93 | 1.31E-06 |
| 1ceyA | 1ceyA | 100 | 128 | 0 | 1 | 128 | 1 | 128 | 7.95E-93 |
| 1ey0A | 1ey0A | 100 | 136 | 0 | 1 | 136 | 1 | 136 | 9.12E-102 |
| 1ihbA | 1ihbA | 100 | 156 | 0 | 1 | 156 | 1 | 156 | 3.99E-115 |
| 1ihbA | 1a5eA | 39.9 | 133 | 79 | 5 | 136 | 16 | 148 | 2.39E-26 |
| 1iobA | 2nvhA | 100 | 152 | 0 | 1 | 152 | 1 | 152 | 3.95E-115 |
| 1l63A | 2lzmA | 98.8 | 162 | 2 | 1 | 162 | 1 | 162 | 3.48E-121 |
| 1lz1A | 1lz1A | 100 | 130 | 0 | 1 | 130 | 1 | 130 | 1.74E-97 |
| 1lz1A | 4lyzA | 60.9 | 128 | 49 | 1 | 128 | 1 | 127 | 1.03E-56 |
| 1lz1A | 1hfzA | 39.3 | 117 | 66 | 3 | 119 | 4 | 115 | 9.82E-30 |
| 1lz1A | 1hmkA | 39.3 | 117 | 66 | 3 | 119 | 4 | 115 | 5.41E-29 |
| 1oh0A | 1oh0A | 100 | 125 | 0 | 1 | 125 | 1 | 125 | 1.42E-94 |
| 1rn1C | 1rn1C | 100 | 104 | 0 | 1 | 104 | 1 | 104 | 2.72E-76 |
| 1vqbA | 1vqbA | 100 | 86 | 0 | 1 | 86 | 1 | 86 | 4.14E-63 |
| 2lzmA | 2lzmA | 100 | 164 | 0 | 1 | 164 | 1 | 164 | 5.50E-125 |
| 2rn2A | 2rn2A | 100 | 155 | 0 | 1 | 155 | 1 | 155 | 1.18E-119 |
| 4lyzA | 4lyzA | 100 | 129 | 0 | 1 | 129 | 1 | 129 | 4.35E-96 |
| 4lyzA | 1lz1A | 60.9 | 128 | 49 | 1 | 127 | 1 | 128 | 1.02E-56 |
| 4lyzA | 1hmkA | 44.2 | 113 | 59 | 3 | 115 | 4 | 112 | 1.61E-30 |
| 4lyzA | 1hfzA | 41.6 | 113 | 62 | 3 | 115 | 4 | 112 | 5.78E-28 |
| 5ptiA | 5ptiA | 100 | 58 | 0 | 1 | 58 | 1 | 58 | 1.85E-41 |

Query represents proteins in the S^sym^ data set; Subject represents proteins in the S2648 data set; %ID is the percent identity of the alignment between the query sequence and the subject sequence; Query start/end and Subject end/end denote the starting and ending positions of the alignment in the query and subject sequences, respectively.

Table S4. Proteins in the VariBench data set (Subject) that are either identical or likely to be homologous to proteins in the S^sym^ data set (Query).

| Query | Subject | %ID | Length | # Mismatches | Query start | Query end | Subject start | Subject end | E-value |
| --- | --- | --- | --- | --- | --- | --- | --- | --- | --- |
| 1bniA | 1bniA | 100 | 108 | 0 | 1 | 108 | 1 | 108 | 1.10E-80 |
| 1bniA | 1mgrA | 37.3 | 59 | 35 | 51 | 107 | 35 | 93 | 7.30E-07 |
| 1bniA | 1rggA | 33.8 | 80 | 44 | 31 | 107 | 19 | 92 | 6.51E-04 |
| 1ey0A | 1stnA | 100 | 136 | 0 | 1 | 136 | 1 | 136 | 5.08E-102 |
| 1iobA | 1iobA | 100 | 153 | 0 | 1 | 153 | 1 | 153 | 5.01E-116 |
| 1l63A | 1l63A | 100 | 162 | 0 | 1 | 162 | 1 | 162 | 1.75E-122 |
| 1l63A | 2lzmA | 98.8 | 162 | 2 | 1 | 162 | 1 | 162 | 1.94E-121 |
| 1lz1A | 1lz1A | 100 | 130 | 0 | 1 | 130 | 1 | 130 | 9.70E-98 |
| 1lz1A | 4lyzA | 60.9 | 128 | 49 | 1 | 128 | 1 | 127 | 5.73E-57 |
| 1lz1A | 1el1A | 52.3 | 130 | 61 | 1 | 130 | 2 | 130 | 5.53E-53 |
| 1lz1A | 1hfzA | 39.3 | 117 | 66 | 3 | 119 | 4 | 115 | 5.47E-30 |
| 1lz1A | 1hfyA | 39.3 | 117 | 66 | 3 | 119 | 3 | 114 | 3.31E-29 |
| 1rn1C | 1rn1B | 100 | 104 | 0 | 1 | 104 | 1 | 104 | 1.51E-76 |
| 1vqbA | 1vqbA | 100 | 86 | 0 | 1 | 86 | 1 | 86 | 2.31E-63 |
| 2lzmA | 2lzmA | 100 | 164 | 0 | 1 | 164 | 1 | 164 | 3.07E-125 |
| 2lzmA | 1l63A | 98.8 | 162 | 2 | 1 | 162 | 1 | 162 | 1.96E-121 |
| 2rn2A | 2rn2A | 100 | 155 | 0 | 1 | 155 | 1 | 155 | 6.59E-120 |
| 4lyzA | 4lyzA | 100 | 129 | 0 | 1 | 129 | 1 | 129 | 2.42E-96 |
| 4lyzA | 1lz1A | 60.9 | 128 | 49 | 1 | 127 | 1 | 128 | 5.68E-57 |
| 4lyzA | 1el1A | 53.1 | 130 | 59 | 1 | 129 | 2 | 130 | 1.51E-50 |
| 4lyzA | 1hfyA | 44.2 | 113 | 59 | 3 | 115 | 3 | 111 | 8.70E-31 |
| 4lyzA | 1hfzA | 41.6 | 113 | 62 | 3 | 115 | 4 | 112 | 3.22E-28 |
| 5ptiA | 1bpiA | 100 | 58 | 0 | 1 | 58 | 1 | 58 | 1.03E-41 |

Query represents proteins in the S^sym^ data set; Subject represents proteins in the VariBench data set; %ID is the percent identity of the alignment between the query sequence and the subject sequence; Query start/end and Subject end/end denote the starting and ending positions of the alignment in the query and subject sequences, respectively.

Table S5. Proteins in the Q3421 data set (Subject) that are either identical or likely to be homologous to proteins in the S^sym^ data set (Query).

| Query | Subject | %ID | Length | # Mismatches | Query start | Query end | Subject start | Subject end | E-value |
| --- | --- | --- | --- | --- | --- | --- | --- | --- | --- |
| 1amqA | 1amqA | 100 | 396 | 0 | 1 | 396 | 1 | 396 | 0 |
| 1bniA | 1bniA | 100 | 108 | 0 | 1 | 108 | 1 | 108 | 2.21E-80 |
| 1ceyA | 1ceyA | 100 | 128 | 0 | 1 | 128 | 1 | 128 | 8.93E-93 |
| 1ey0A | 1stnA | 100 | 136 | 0 | 1 | 136 | 1 | 136 | 1.03E-101 |
| 1ihbA | 1ihbA | 100 | 156 | 0 | 1 | 156 | 1 | 156 | 4.49E-115 |
| 1ihbA | 1a5eA | 39.9 | 133 | 79 | 5 | 136 | 16 | 148 | 2.69E-26 |
| 1iobA | 1iobA | 100 | 153 | 0 | 1 | 153 | 1 | 153 | 1.01E-115 |
| 1l63A | 1l63A | 100 | 162 | 0 | 1 | 162 | 1 | 162 | 3.53E-122 |
| 1l63A | 2lzmA | 98.8 | 162 | 2 | 1 | 162 | 1 | 162 | 3.92E-121 |
| 1lz1A | 1lz1A | 100 | 130 | 0 | 1 | 130 | 1 | 130 | 1.96E-97 |
| 1lz1A | 4lyzA | 60.9 | 128 | 49 | 1 | 128 | 1 | 127 | 1.16E-56 |
| 1lz1A | 1el1A | 52.3 | 130 | 61 | 1 | 130 | 2 | 130 | 1.12E-52 |
| 1lz1A | 1hfyA | 39.3 | 117 | 66 | 3 | 119 | 3 | 114 | 6.67E-29 |
| 1oh0A | 1oh0A | 100 | 125 | 0 | 1 | 125 | 1 | 125 | 1.6E-94 |
| 1rn1C | 1rn1A | 99.0 | 104 | 0 | 1 | 104 | 1 | 103 | 1.16E-73 |
| 1vqbA | 1vqbA | 100 | 86 | 0 | 1 | 86 | 1 | 86 | 4.66E-63 |
| 2lzmA | 2lzmA | 100 | 164 | 0 | 1 | 164 | 1 | 164 | 6.19E-125 |
| 2lzmA | 1l63A | 98.8 | 162 | 2 | 1 | 162 | 1 | 162 | 3.96E-121 |
| 2rn2A | 2rn2A | 100 | 155 | 0 | 1 | 155 | 1 | 155 | 1.33E-119 |
| 4lyzA | 4lyzA | 100 | 129 | 0 | 1 | 129 | 1 | 129 | 4.89E-96 |
| 4lyzA | 1lz1A | 60.9 | 128 | 49 | 1 | 127 | 1 | 128 | 1.15E-56 |
| 4lyzA | 1el1A | 53.0 | 130 | 59 | 1 | 129 | 2 | 130 | 3.05E-50 |
| 4lyzA | 1hfyA | 44.2 | 113 | 59 | 3 | 115 | 3 | 111 | 1.76E-30 |
| 5ptiA | 1bpiA | 100 | 58 | 0 | 1 | 58 | 1 | 58 | 2.08E-41 |

Query represents proteins in the S^sym^ data set; Subject represents proteins in the Q3421 data set; %ID is the percent identity of the alignment between the query sequence and the subject sequence; Query start/end and Subject end/end denote the starting and ending positions of the alignment in the query and subject sequences, respectively.

Table S6. A brief summary of the characteristics of methods presented in Table 2.

| Method | Algorithm | Feature | Reference |
| --- | --- | --- | --- |
| DDGun3D | Linear parametric model | BLOSUM62 substitution matrix, statistical potentials, hydrophobicity, solvent accessibility, and evolutionary information derived from multiple sequence alignment | (Montanucci et al., 2019) |
| DDGun | Linear parametric model | BLOSUM62 substitution matrix, statistical potentials, hydrophobicity, and evolutionary information derived from multiple sequence alignment | (Montanucci et al., 2019) |
| PoPMuSiC^sym^ | Same as PoPMuSiC 2.1 except two coefficients are constrained | Statistical potentials, amino acid volume | (Pucci et al., 2015) |
| MAESTRO | Linear regression, neural network, and support vector machine | Statistical potentials | (Laimer et al., 2016) |
| FoldX | NA | Empirical force field | (Guerois et al., 2002) |
| PoPMuSiC 2.1 | Linear parametric model with coefficients fitted by a neural network | Statistical potentials, amino acid volume | (Dehouck et al., 2009) |
| SDM | NA | Environment-specific amino acid substitution table | (Worth et al., 2011) |
| iSTABLE | Meta predictor | Evolutionary information and predictions from I-Mutant, AUTOMUTE, MUPRO, PoPMuSiC, and CUPSAT | (Chen et al., 2013) |
| I-Mutant 3.0 | Support vector machine | Protein sequence and structure information | (Capriotti et al., 2005) |
| NeEMO | Neural network | Features extracted from residue interaction networks | (Giollo et al., 2014) |
| DUET | Support vector machine | Predictions from mCSM and SDM | (Pires et al., 2014a) |
| mCSM | Gaussian progress regression and random forest | Graph-based distance patterns | (Pires et al., 2014b) |
| MUPRO | Support vector machine | Sequence information | (Cheng et al., 2006) |
| STRUM | Gradient boosting regression | Sequence information, structure information, and evolutionary information derived from multiple sequence alignment | (Quan et al., 2016) |
| Rosetta | NA | Empirical energy function | (Simons et al., 1997) |
| AUTOMUTE | Support vector machine and random forest | Statistical potentials | (Masso and Vaisman, 2008) |
| CUPSAT | NA | Statistical potentials | (Parthiban et al., 2006) |

NA indicates either no machine-learning algorithm or no statistical model was used.

Table S7. Comparison of ThermoNet with four other methods on p53.

| Method | $\boldsymbol{\sigma}_{\boldsymbol{dir}}$ | $\mathbf{r}_{\boldsymbol{dir}}$ | $\boldsymbol{\sigma}_{\boldsymbol{rev}}$ | $\mathbf{r}_{\boldsymbol{rev}}$ | $\mathbf{r}_{\boldsymbol{dir-rev}}$ | $\left\langle\boldsymbol{\delta} \right\rangle$ |
| --- | --- | --- | --- | --- | --- | --- |
| FoldX | 1.79 | 0.71 | 1.51 | 0.75 | -0.97 | 0.23 |
| ThermoNet | 2.01 | 0.45 | 1.92 | 0.56 | -0.93 | -0.04 |
| Rosetta | 3.47 | 0.78 | 2.75 | 0.70 | -0.84 | 1.15 |
| SDM | 1.52 | 0.68 | 2.44 | 0.01 | -0.37 | 0.71 |
| CUPSAT | 2.83 | 0.25 | NA | NA | NA | NA |

Table S8. Comparison of ThermoNet with four other methods on myoglobin.

| Method | $\boldsymbol{\sigma}_{\boldsymbol{dir}}$ | $\mathbf{r}_{\boldsymbol{dir}}$ | $\boldsymbol{\sigma}_{\boldsymbol{rev}}$ | $\mathbf{r}_{\boldsymbol{rev}}$ | $\mathbf{r}_{\boldsymbol{dir-rev}}$ | $\left\langle\boldsymbol{\delta} \right\rangle$ |
| --- | --- | --- | --- | --- | --- | --- |
| FoldX | 1.40 | 0.61 | 1.36 | 0.60 | -0.97 | 0.12 |
| ThermoNet | 1.16 | 0.38 | 1.18 | 0.37 | -0.97 | -0.02 |
| Rosetta | 4.72 | 0.63 | 3.62 | 0.63 | -0.93 | 0.91 |
| SDM | 1.25 | 0.52 | 1.48 | 0.12 | 0.18 | 0.89 |
| CUPSAT | 1.60 | 0.25 | NA | NA | NA | NA |

NA: not available. The CUPSAT server does not offer batch processing for predicting the ∆∆Gs of reverse mutations nor a downloadable standalone version for local use.

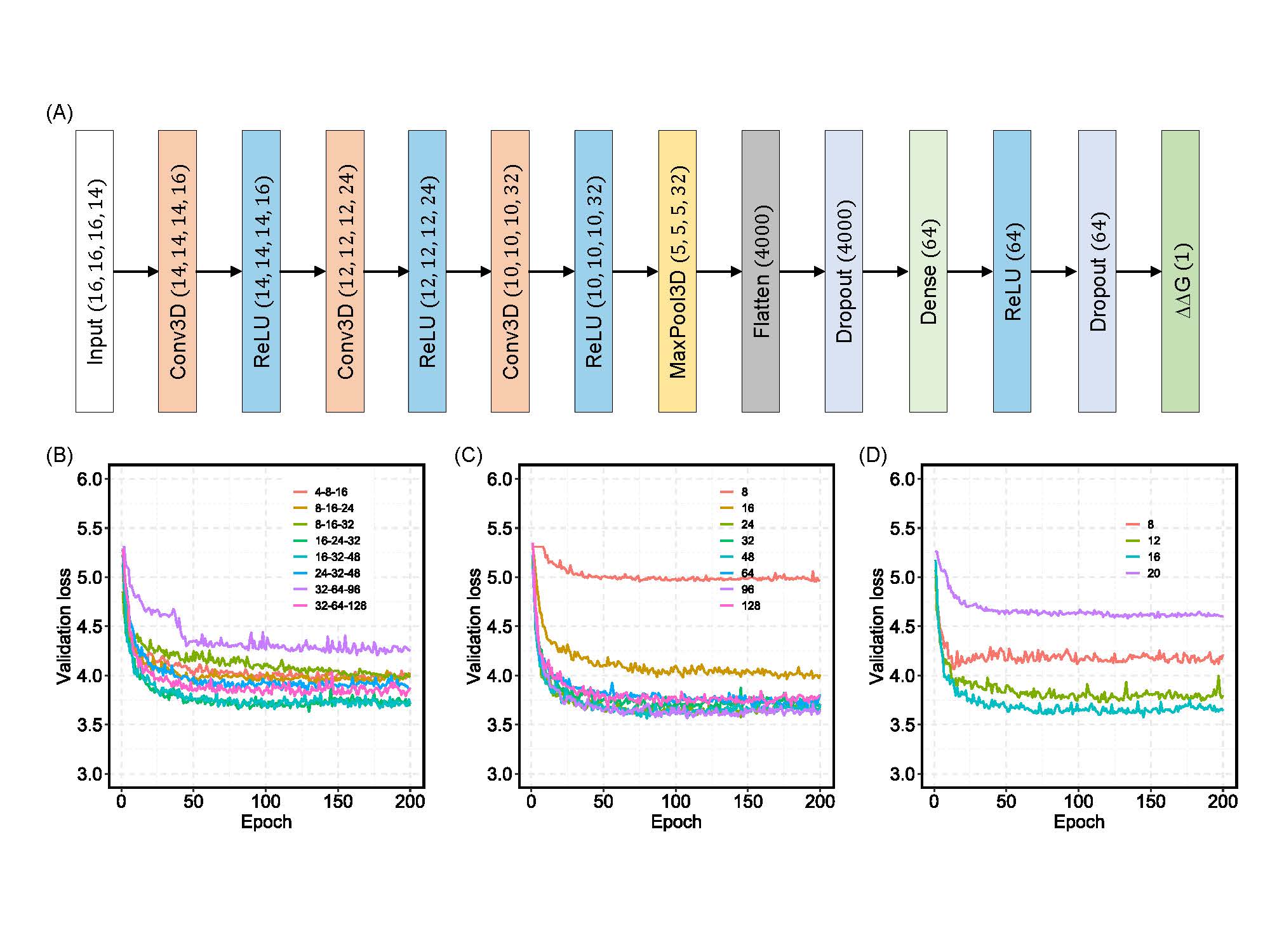

Fig S1. Architecture of the deep 3D convolutional neural network model and hyperparameter search.

(A) The overall organization of the model begins with the input tensor and ends with a final layer that outputs ∆∆G prediction. The numbers in parentheses before the Flatten layer represent the dimensionality of the output from each layer in the format (width, height, depth, property channels). The number in parentheses starting from the Flatten layer represent the number of output features from each of the densely connected layers. This optimized architecture was determined through cross-validation. (B) Results from cross validating the sizes of the convolutional layers while keeping the size of the densely connected layer at 32 neurons. (C) Results from cross validating the size of the densely connected layer while keeping the sizes of the convolutional layers at (16, 24, 32). (D) Results from cross validating the dimensions of the input grid.

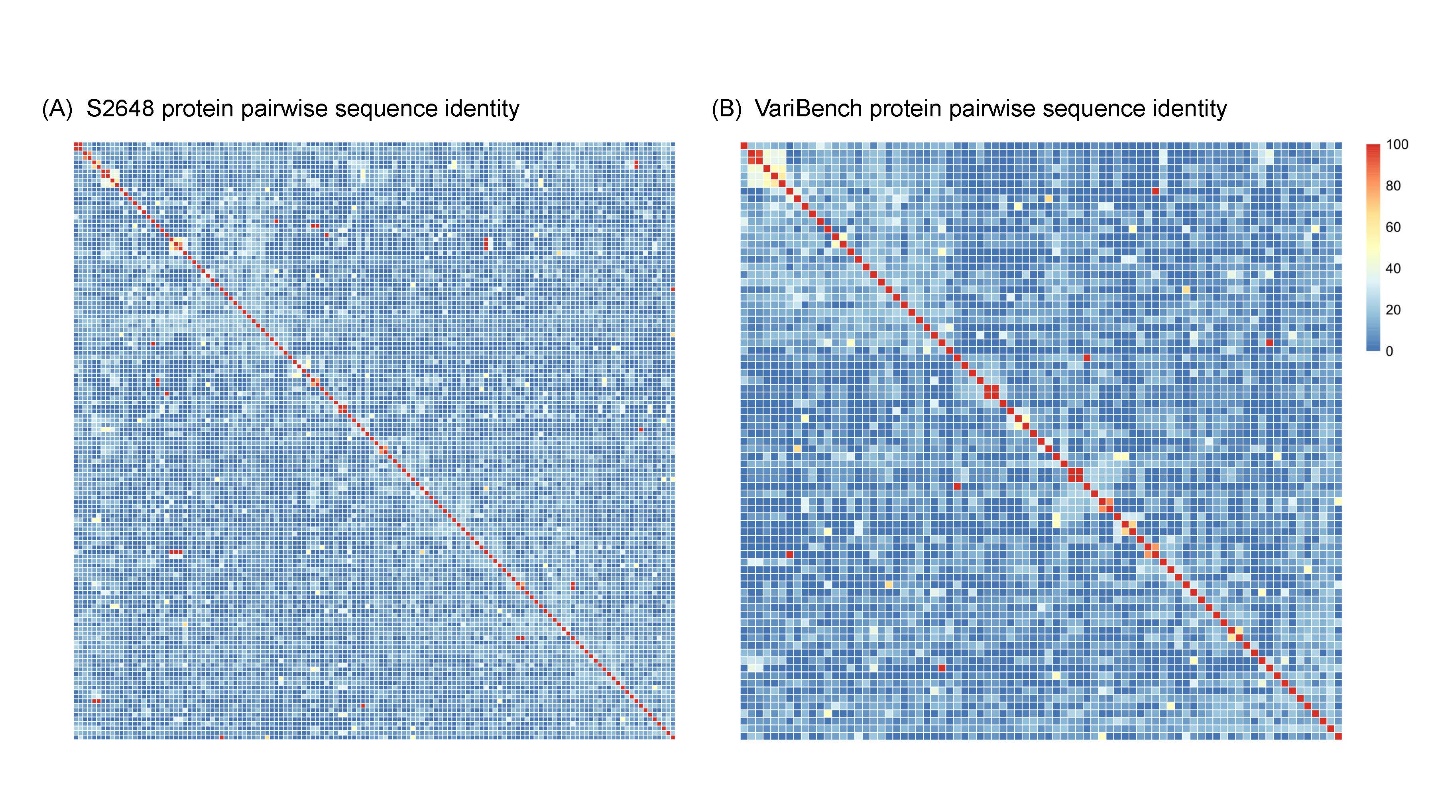

Fig S2. Pairwise percent sequence identity of the proteins in the S2648 and VariBench data sets. (A) A heatmap representation of the pairwise percent sequence identity matrix of the proteins in the S2648 data set. (B) A heatmap representation of the pairwise percent sequence identity matrix of the proteins in the VariBench data set. It is obvious from these two heatmaps that there is substantial pairwise homology (percent identity > 25%) in both S2648 and VariBench. The pairwise identity matrices were obtained using the Clustal Omega multiple sequence alignment program (Sievers et al., 2011).

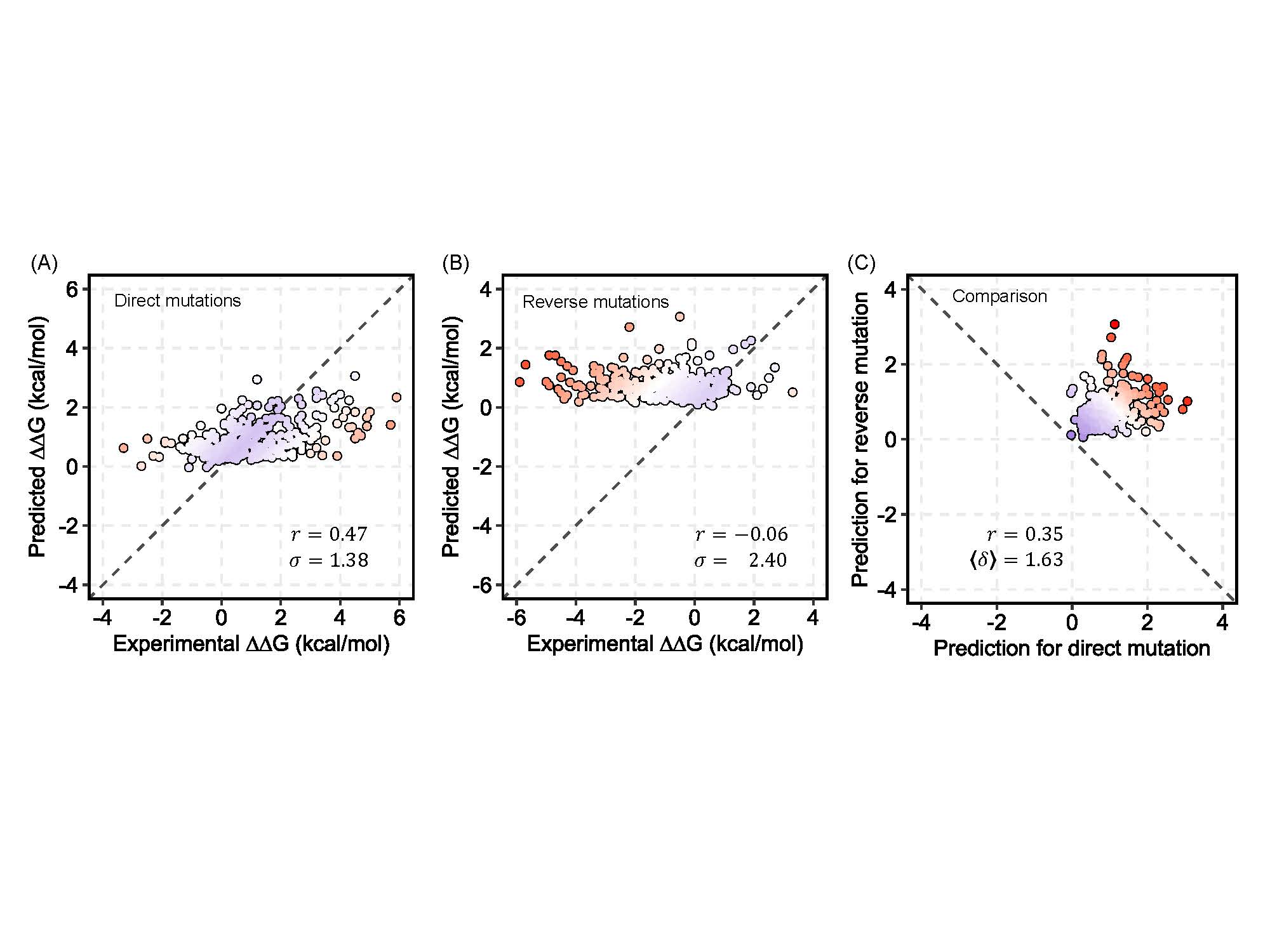

Fig S3. CNNs trained using only direct mutations show a large prediction bias.

(A) Performance of an ensemble of ten networks trained using only the set of 1,744 direct mutations on predicting the ∆∆Gs of the direct mutations in the blind test set; The Pearson correlation coefficient ($r$) between predicted values and experimentally determined values is 0.47, and the root-mean-square deviation ($\sigma$) of predicted values from experimentally determined values is 1.38 kcal/mol. (B) Performance of the same ensemble of ten networks on predicting the ∆∆Gs of the reverse mutations in the blind test set; The Pearson correlation coefficient ($r$) between predicted values and experimentally determined values is -0.06, and the root-mean-square deviation ($\sigma$) of predicted values from experimentally determined values is 2.40 kcal/mol. (C) Direct versus reverse ∆∆G values of all the mutations in the blind test set predicted by the same ensemble of networks. (B) and (C) highlight that the models trained with only direct mutations have a large bias and, when compared to the models trained using the balanced data set, the necessity of adding reverse mutations to correct the bias. The dots are colored in gradient from blue to red such that blue represents the most accurate prediction and red represents the least accurate.
